# Balancing activation and costimulation of CAR tunes signaling dynamics and enhances therapeutic potency

**DOI:** 10.1101/2022.03.01.482445

**Authors:** Yanting Duan, Jiangqing Chen, Xianhui Meng, Longwei Liu, Kai Shang, Xiaoyan Wu, Yajie Wang, Zihan Huang, Houyu Liu, Yanjie Huang, Chun Zhou, Xiaofei Gao, Yingxiao Wang, Michel Sadelain, Jie Sun

## Abstract

**Background:** Primary human T cells engineered with chimeric antigen receptors (CARs) ex vivo can be adoptively transferred to treat cancer. CD19-targeting CAR with CD28 costimulatory domain and CD3ζ activation domain have been approved by the US FDA for treating B cell malignancies.

**Methods:** Here we generated mutation of immunorecpetor tyrosine-based activation motifs (ITAMs) in CD3ζ, namely 1XX CAR, which altered the balance of activation and costimulation. Next we investigated whether 1XX design could enhance therapeutic potency against solid tumors. We constructed both CD19- and AXL-specific 1XX CARs and compared their *in vitro* and *in vivo* functions with their WT counterparts.

**Results:** Even though 1XX CARs decreased cytotoxicity against tumor cells *in vitro*, they showed better anti-tumor efficacy in both pancreatic and melanoma mouse models. Detailed analysis revealed that 1XX CAR-T cells proliferated more in response to antigen stimulation *in vitro*, persisted longer *in vivo* and had higher percentage of central memory cells. As 1XX modification directly calibrates CAR activation potential, we utilized fluorescence resonance energy transfer (FRET)-based biosensor to monitor signaling dynamics downstream of CARs. Decreased ITAM numbers in 1XX resulted in similar ZAP70 activation, while 1XX induced higher Ca^2+^ elevation and faster Erk activation than WT CAR, which may contribute to the better therapeutic potency of 1XX.

**Conclusions:** Our results established the surpiosity of 1XX against two targets in different solid tumor models and shed light on the underlying molecular mechanism of CAR signaling, paving the way for the clinical application of 1XX CARs against solid tumors.

## Background

The adoptive transfer of chimeric antigen receptor (CAR)-T cells has achieved great success in treating hematological maliganancies and the US FDA has approved four CD19-specific CAR-T products and one product targeting BCMA ^[1]^. On the other hand, it is generally recognized that the application of CAR-T in solid tumors has been impeded by many factors, including nonspecific tumor antigen, inefficient tumor trafficking and immunosuppressive tumor microenvironment ^[2]^. Various strategies have been developed in combination of CAR to overcome these hurdles, such as cytokines ^[3,4]^, scFv against PD-1 ^[5]^, BiTEs to recruit bystander T cells ^[6]^ and the use of IL-7/CCL-19 to increase trafficking ^[7]^. However, the CAR design itself may still not be optimized to fully harness the power of T cells.

In addition to the activation signal from T cell receptor (TCR), T cells integrate signals from costimulatory/coinhibitory receptors and cytokine receptors though a complex signaling network to direct multiple functions, such as proliferation, cytotoxicity, cytokine secretion and differentiation ^[8]^. These functions need to be balanced to achieve optimal long-term anti-tumor effects. After antigen binding, CD28-based second-generation CARs transduce signals through costimulatory domain from CD28 and activation domain from CD3ζ, which comprises three immunorecpetoro tyrosine-based activation motifs (ITAMs). Both pathways highly depend on the activity of Lck kinase. Each ITAM in CD3ζ after phosphorylated by Lck recruits ZAP70 kinase so that ZAP70 can be phosphorylated by Lck ^[9]^. ZAP70 activation further phosphorylates adaptor LAT at the membrane, which forms a signaling network with various proteins, such as Grb2/Sos and PLCγ. PLCγ catalyzes the formation of two important second messengers, DAG and IP3 ^[10]^. IP3 binds to receptors at ER to release large amount of Ca^2+^, while both DAG and Grb2/Sos could activate Erk through Ras ^[11,12]^. Meanwhile, CD28 cytosolic domain after being phosphorylated by Lck can recruit and activate PI3K ^[13]^. PIP3 produced by PI3K binds to PH domain of Itk, recruiting Itk to the membrane. Itk can directly interact with PLCγ and activate it ^[14]^. Thus both pathways can modulate downstram Ca^2+^ and Erk activities.

A single ITAM-containing 1XX CAR design with calibrated activation potential has been shown to have superior anti-tumor activity in a leukemia mouse model by balancing activation and costimulatory signals ^[15]^. Here we investigated whether this design benefits CAR-T therapy against solid tumors. We also quantified the molecular activities in CAR-T cells with high temporal resolution by fluorescence resonance energy transfer (FRET) biosensors. Based on the importance of the signaling molecule and the avaibility of biosensors, we chose ZAP70 ^[16]^, Ca^2+ [17]^ and Erk ^[18]^ biosensors to reveal differences in signaling dynamics induced by 1XX compared to WT CAR.

## Results

### CD19-1XX CAR-T cells had lower cytotoxicity against Panc-1 cells *in vitro*

In the NALM6 leukemia mouse model, CD19-1XX CAR-T cells had shown superior anti-tumor activity than CD19-WT cells. In this study, we first investigated whether the CD19-1XX CAR-T cells control solid tumors better than CD19-WT cells. With CRISPR/Cas9-mediated gene targeting ^[19]^, we integrated CD19-WT or CD19-1XX CAR at the specific *TRAC* locus to generate CAR-T cells (Fig. 1a, Additional file 1: Fig. 1a). The resulting CAR-T cells have similar knock-in efficiency (Fig. 1b) and surface expression of CARs (Fig. 1c). To have a direct comparison of the two CD19-specific CAR-T cells, we ectopically expressed the human CD19 antigen in the pancreatic cell line Panc-1for both *in vitro* killing assay and *in vivo* xenograft model (Additional file 1: Fig. 1b). With Panc-1^CD19+^ cells as targets, CD19-1XX CAR-T cells displayed significantly lower cytotoxicity than CD19-WT cells at different E:T ratios in an 18h killing assay (Fig. 1d, e). 24h after antigen stimulation, CD19-1XX and CD19-WT CAR-T cells secreted similar amount of IL-2 and TNFα (Fig. 1f, g), while IFNγ secreted by CD19-1XX was about 15% lower than that secreted by CD19-WT (Fig. 1f, g). In addition, our CFSE-based assay showed that 72h after antigen stimulation, CD19-1XX cells had 8 fold increase of proliferation while CD19-WT cells only had 1.7 fold increase (Fig. 1h, i). This was mainly attributed to the much lower baseline proliferation of 1XX cells in the absence of antigen (Fig. 1h), suggesting 1XX had lower tonic signaling. In summary, CD19-1XX cells displayed lower cytotoxicity and IFN γ secretion than CD19-WT ones while they showed higher antigen-dependent proliferation *in vitro*.

**Fig.1.**
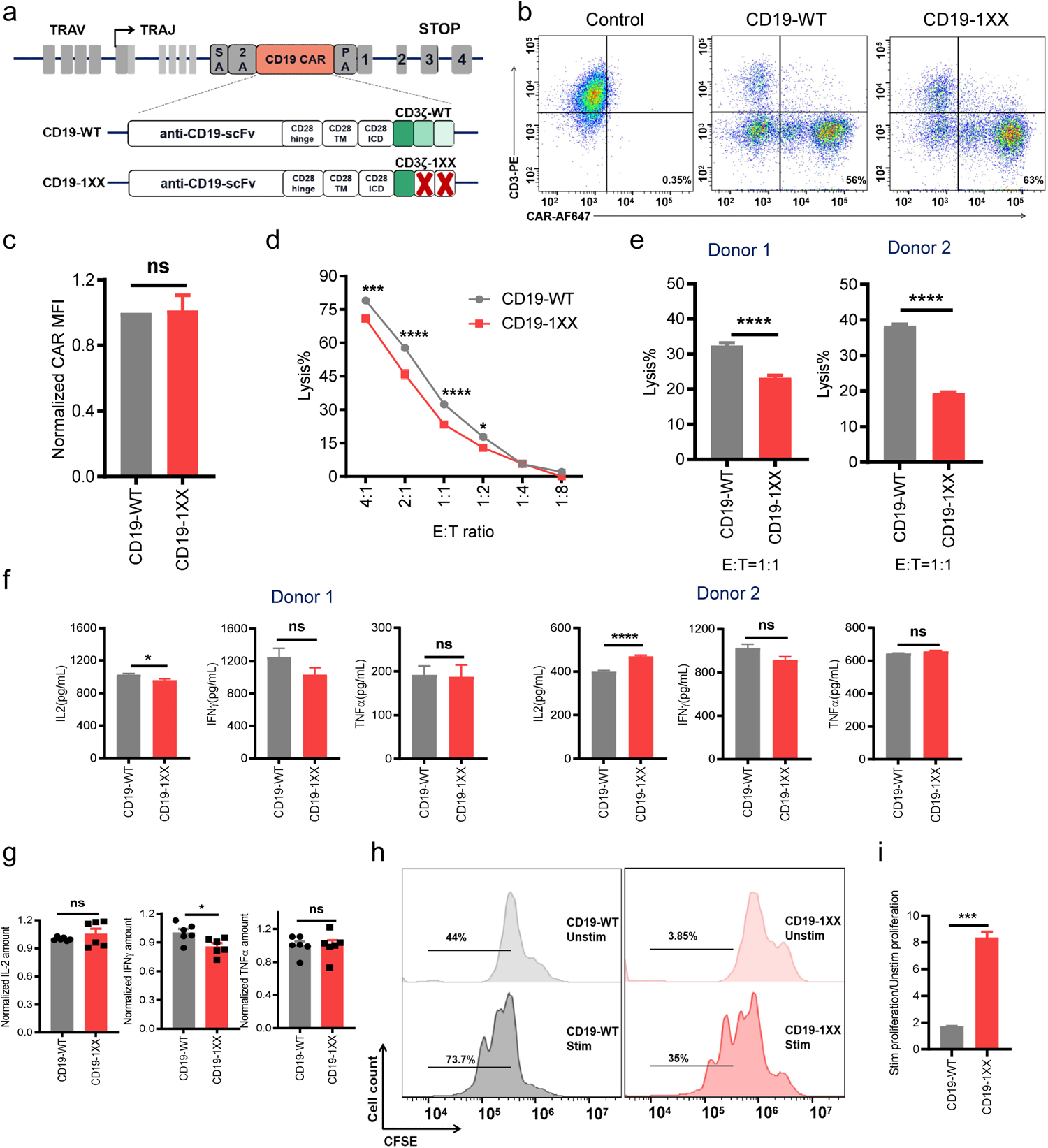
CD19-1XX CAR-T cells show reduced cytotoxicity but enhanced proliferation in vitro. (a) Targeting of CD19-WT or CD19-1XX CAR into the *TRAC* locus by CRISPR/Cas9. (b)FACS analysis showing the expression levels of two CD19 CARs. Data are representative of at least three independent experiments with similar results. Untransduced T cells were used as the control. (c) Normalized CD19 WT and 1XX CAR expression (n=3). (d) Cytotoxic activity of CD19-WT and CD19-1XX CAR-T cells using an 18 h bioluminescence assay with FFL-expressing Panc-1^CD19+^ cells as targets. (n=3, 1 experiment performed in triplicates from 1 donor). (e) The lysis of target cells by CD19-WT or CD19-1XX CAR-T cells at E:T=1:1 (n=6, 2 independent experiments performed in triplicates from 2 donors). (f,g) Cytokine secretion of CD19-WT and CD19-1XX cells were detected after stimulation by target cells. (E:T=4:1, n=6, 2 independent experiments performed in triplicates from 2 donors). (h) The proliferation of CD19-WT and CD19-1XX CAR-T cells analyzed by CSFE assay. Unstim represents CAR-T cells without target cell coculture and Stim represents CAR-T cells stimulated with target cells for 72h. (E:T=4:1, n=3, 1 experiment performed in triplicates from 1 donor). (i) CAR-T cell antigen-dependent proliferation fold caculated by Stim/Unstim ratio.

### CD19-1XX CAR-T cells showed better anti-tumor activity in a pancreatic tumor model

We next compared the anti-tumor activity of two CD19 CAR-T cells in a xenograft pancreatic cancer model, where Panc-1^CD19+^ cells were subcutaneously injected in the right flank of NSG mice (Fig. 2a). We used a low dose of CAR-T cells (1×10^6^) to better compare T cell potency in the CAR stress test ^[20]^. Though both CAR-T cells inhibited initial tumor growth, mice injected with CD19-WT CAR-T cells started to relapse around day 29 while mice treated with CD19-1XX CAR-T cells relapsed around day 46 (Fig. 2b). At day 53, the average tumor volume of mice with CD19-1XX CAR-T cells was significantly smaller than that of mice injected with CD19-WT cells (Fig. 2b, c). As CAR targeting at *TRAC* locus knocked out CD3 expression, we used antibodies against human CD4 or CD8 molecule to stain for human CAR-T cells in the tumor tissue. Our results revealed the existence of CD4^+^ and CD8^+^ T cells in the tissue even at day 53, albeit at a very low percentage (Fig. 2d-f). Even though the percentage of either CD4^+^ or CD8^+^ T cells was not statistically different between CD19-1XX and CD19-WT groups, CD19-1XX groups has a trend of more CD4^+^ and CD8^+^ T cells (Fig. 2e, f).

**Fig.2.**
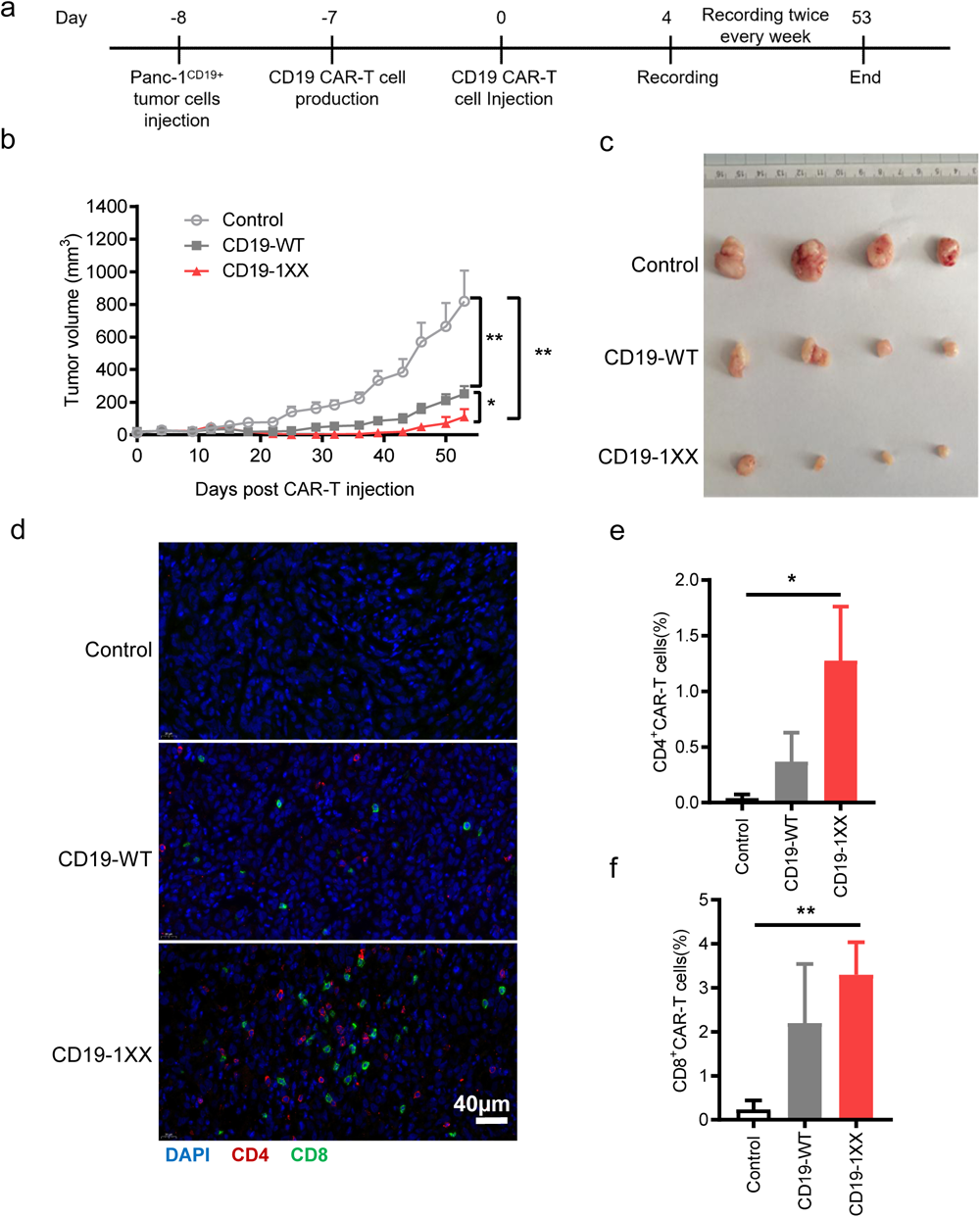
CD19-1XX CAR-T cells show better anti-tumor activity in a pancreatic tumor model in vivo. (a) Timeline of of CD19 CAR-T treatment in mouse model. (b) Panc-1^CD19+^ -bearing mice were treated with 1 × 10^6^ CD19 CAR-T cells as indicated and tumor burden (tumor volume) of mice was measured at indicated days (n=4 mice each group). For mouse experiments, Control, PBS. (c) Tumor tissue from mice treated with indicated CAR-T cells. (d) Tumor tissue from mice treated with indicated CAR-T cells was sliced and stained with antibodies against human CD4 and CD8 (n=4). The percentage of CD4^+^ (e) and CD8^+^ (f) T cells in tumor tissue from different groups of mice (n=4). All data are mean ± SEM.

### 1XX modification of AXL CAR enhanced its anti-tumor activity in a melanoma model

After we confirmed that CD19-1XX cells were better than CD19-WT in both leukemia and solid tumor models, we further investigated whether 1XX design is also beneficial for CARs targeting antigens expressed naturally in solid tumors. AXL is a tyrosine kinase receptor, commonly overexpressed in both hematological malignancies and solid tumors ^[21]^. Human melanoma cell line A375 was shown to have a high expression of AXL antigen (Additional file 1: Fig. 2a). We constructed a CAR targeting AXL (AXL-WT) using the scFv from an DAXL-88 antibody we reported before (Additional file 1: Fig. 2b) and also generated 1XX-modified AXL-specific CAR for comparison (Fig. 3a). Expression of AXL-WT and AXL-1XX CAR was also similar (Fig. 3b, c). In terms of cytotoxicity against A375 cell line, AXL-1XX was lower than AXL-WT CAR-T cells (Fig. 3d, e). Meanwhile, data summarized from two donors suggested that AXL-1XX and AXL-WT CAR-T cells had similar cytokine secretion (Fig. 3f, g). CFSE-based assay showed AXL-1XX has stronger proliferative capability than AXL-WT upon antigen-stimulation (Fig. 3h, i).

**Fig.3.**
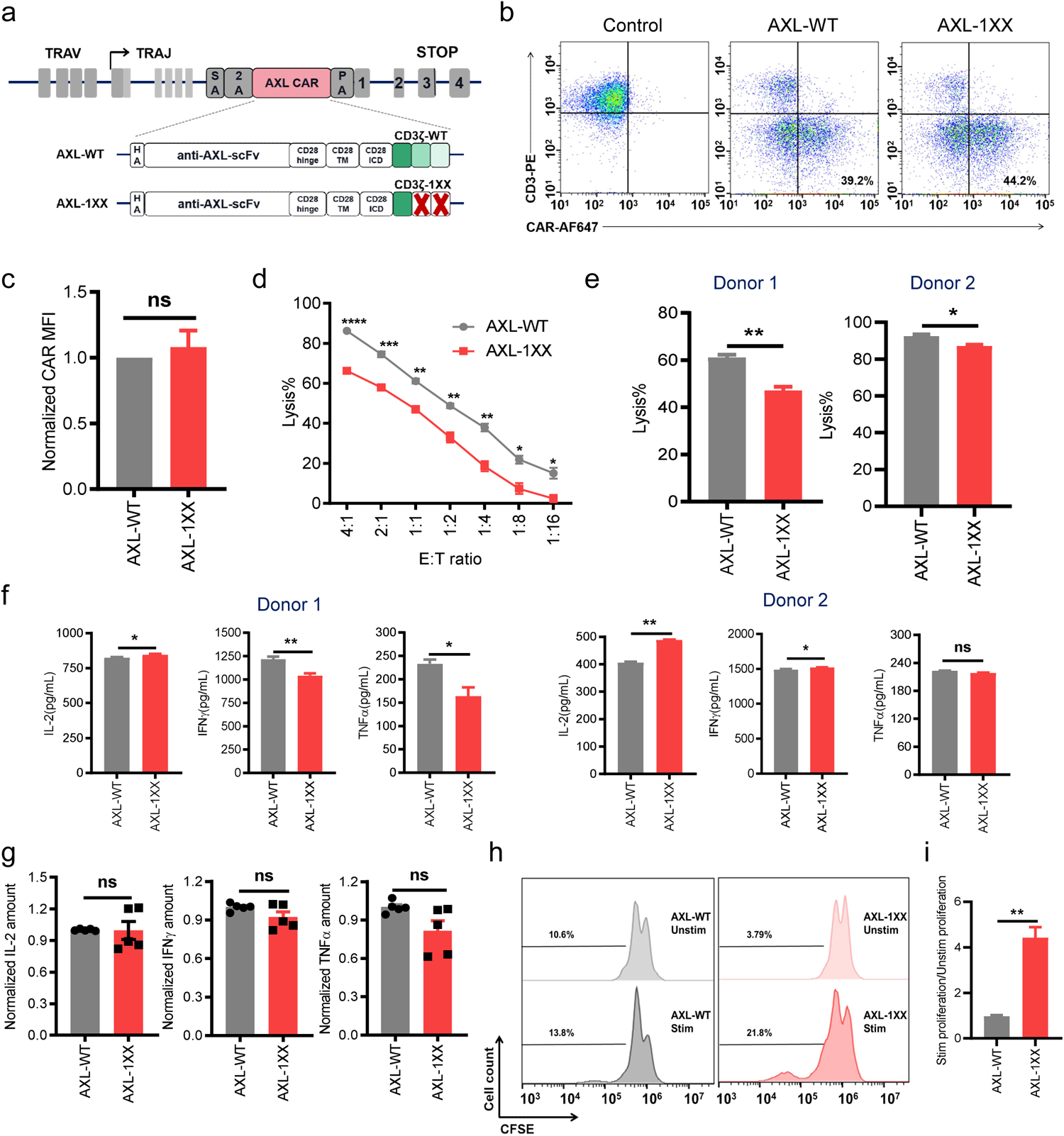
1XX modification of AXL CAR reduced cytotoxicity but enhanced proliferation in vitro. (a) Targeting of AXL-WT or AXL-1XX into the *TRAC* locus by CRISPR/Cas9. (b) FACS analysis showing the expression levels of two AXL CARs. Data are representative of at least three independent experiments with similar results. Untransduced T cells were used as the control. (c) Normalized CD19 WT and 1XX CAR expression (n=3). (d) Cytotoxic activity of AXL-WT and AXL-1XX using an 18 h bioluminescence assay with FFL-expressing A375 cells as targets. (n=3, 1 experiment performed in triplicates from 1 donor). (e) The lysis of target cells by AXL-WT or AXL-1XX CAR-T cells at E:T=1:1 (n=6, 2 independent experiments performed in triplicates from 2 donors). (f,g) Cytokine secretion of AXL-WT and AXL-1XX were detected 24h after stimulation by target cells (E:T=4:1, n=6, 2 independent experiments performed in triplicates from 2 donors). (h)The proliferation of CD19-WT and CD19-1XX CAR-T cells analyzed by CSFE assay. Unstim represents CAR-T cells without target cell coculture and Stim represents CAR-T cells stimulated with target cells for 72h. (E:T=4:1, n=3, 1 experiment performed in triplicates from 1 donor). (i) CAR-T cell antigen-dependent proliferation fold caculated by Stim/Unstim ratio.

Then we compared their anti-tumor activity in a xenograft melanoma model, where A375 cells were subcutaneously injected in the right flank of NSG mice (Fig. 4a). Interestingly, AXL-WT CAR-T failed to control tumor growth when injected at a low 1×10^6^ dose while AXL-1XX CAR-T cells significantly reduced tumor size at the same dose (Fig. 4b-d). The analysis of CAR-T cells extracted from mice revealed that the total number of AXL-1XX CAR-T cells was about 110 fold more than that of AXL-WT cells (Fig. 4e). The dramatic increase of cell number from AXL-1XX CAR-T cells were seen in both CD4^+^ (116 fold) and CD8^+^ pupulations (121 fold) (Fig. 4f, g). Further phenotypic analysis showed that AXL-1XX CAR-T cells had significantly higher percentage of central memory cells and lower percentage of effector cells in both CD4^+^ and CD8^+^ populations (Fig. 4h, i). As total CAR-T cells were much higher in 1XX, the total numbers of central memeory cells and effector cells were both higher in mice injected with AXL-1XX cells (Fig. 4j, k). Meanwhile, 1XX led to the trend of fewer percentage of PD-1^+^LAG3^+^ exhausted cells than AXL-WT cells (Additional file 1: Fig. 2c). To summarize, the comparison of AXL-1XX and AXL-WT was mostly consistent with CD19-1XX and CD19-WT comparison. Although 1XX design reduced CAR-T cytotoxicity *in vitro*, but it enhanced the anti-tumor activity of CARs targeting two solid tumor models *in vivo*, which has more significance for CAR-T therapy.

**Fig.4.**
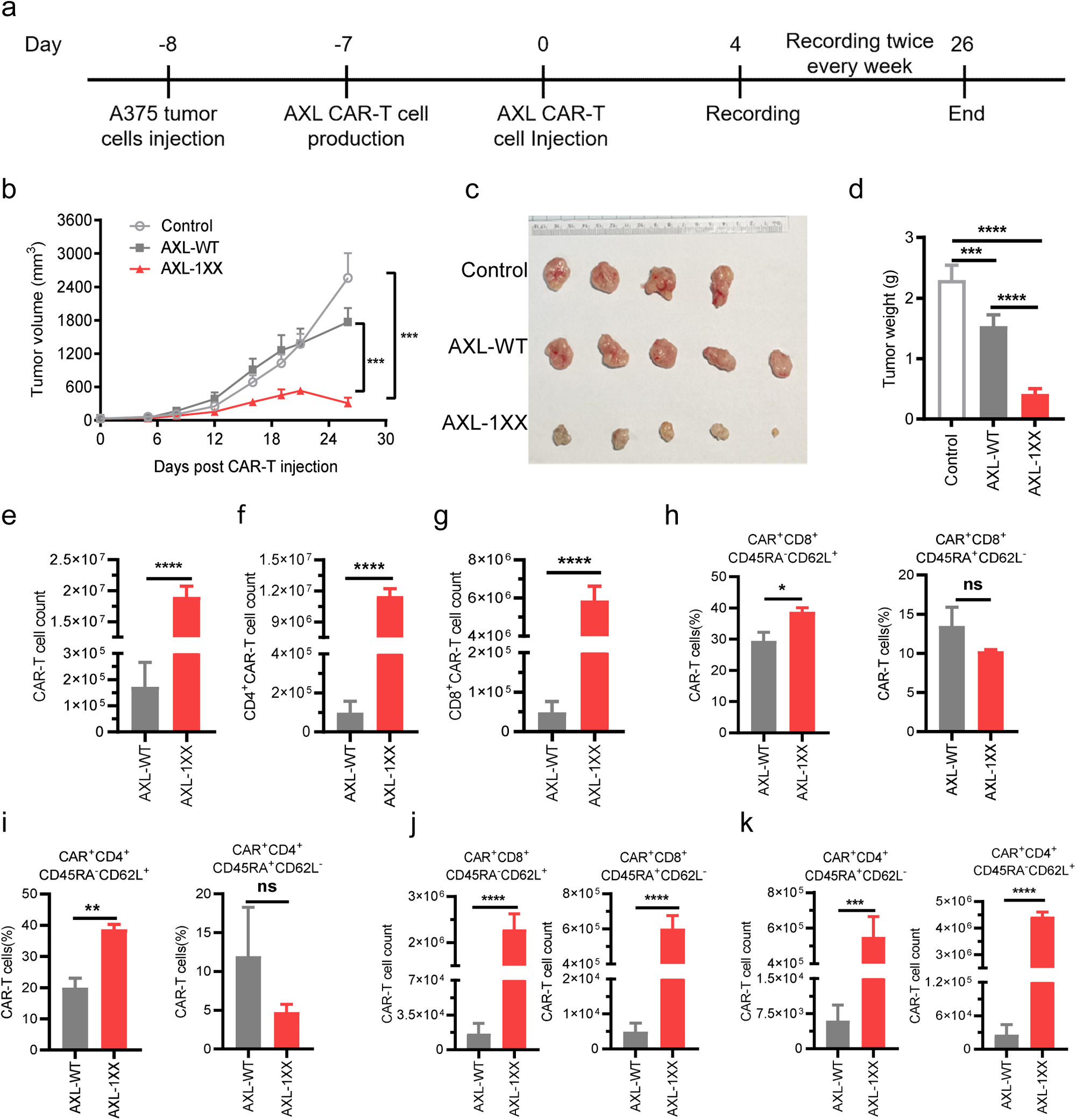
1XX modification of AXL CAR enhance its anti-tumor activity in a melanoma tumor model. (a) Timeline of of AXL CAR-T treatment in mouse model. (b) A375-bearing mice were treated with 2.5 × 10^6^ AXL CAR-T cells and tumor burden (tumor volume) of mice were measured at indicated days (n=4, 5, 5 mice in respective groups). Control, PBS. (c) Tumor tissue of mice treated with indicated CAR-T cells (n=4, 5, 5). (d) The weight of tumors from different groups of mice (n=4, 5, 5). Number of CAR-T(e), CD4^+^ CAR-T cells (f) and CD8^+^ CAR-T cells (g) in spleen of mice 20 days post CAR-T infusion (n=5, 3). (h-k) Phenotype of CAR-T cells in the spleen of mice 20 days after CAR-T infusion as demonstrated by the percentage of T_CM_ (CD45RA^-^CD62L^+^) and T_EFF_ (CD45RA^+^CD62L^-^) cells (n=5, 3). All data are mean ± SEM.

### 1XX CAR induced higher Ca^2+^ level and faster Erk activation than WT CAR

Our previous work in leukemia model indicated that CD19-1XX cells had a better anti-tumor activity because they were more persistent and comprised of increased numbers of memory cells. It is intriguing how a simple 1XX calibration can have profound impact on cellular phenotype and functions. To complement the global RNAseq comparison between CD19-1XX and CD19-WT ^[15]^, we next compared CAR-mediated signaling dynamics in CD19-1XX and CD19-WT cells with different FRET biosensors. FRET biosensors enable monitoring of the molecular activity at single cell level with high spatiotemporal resolution. Here all the FRET changes of biosensor in every cell was normalized to its unstimulated baseline level so we could directly compare antigen-induced activation.

As ZAP70 kinase directly binds phosphorylated ITAMs of TCR and CARs to get activated, 1XX mutation might directly affect the activity of ZAP70. Using a newly developed ZAP70 FRET biosensor with improved dynamic range, ZAP70 activation dynamics can be observed in CAR-T cells upon antigen stimulation ^[16]^. We expressed this biosensor together with either CD19-1XX or CD19-WT CAR and performed live cell FRET imaging (Fig. 5a). ZAP70 activation can be observed by the FRET biosensor after CAR-antigen interaction and the activation dynamics induced by CD19-1XX and CD19-WT CAR were not statistically different (Fig. 5b-e, Additional file 1: Fig. 3a-b). However, CD19-1XX exhibited a trend of lower ZAP70 activity within the first 5min but eventually catched up with CD19-WT at 30min. We next investigated cytosolic Ca^2+^ evelation ^[17]^ and Erk activation, as they are key signaling events downtream of both TCR/CD3ζ-ZAP70 and CD28-PI3K pathways and have profound effects on T cell functions through transcriptional regulation. Our results showed that Ca^2+^ elevation at 1.5min in 1XX cells was significantly higher than that in WT cells (Fig. 5f-j, Additional file 1: Fig. 3c, d). Moreover, analysis of CAR-T cells transduced with Erk biosensors ^[18]^ revealed that CD19-1XX directed faster Erk activation than CD19-WT did upon target cell binding (Fig. 5k-o, Additional file 1: Fig. 3e, f). Thus, even though 1XX-induced a trend of reduced ZAP70 activity, it increased antigen-dependent Ca^2+^ elevation and sped up Erk activation, which were correlated with higher therapeutic potency.

**Fig.5.**
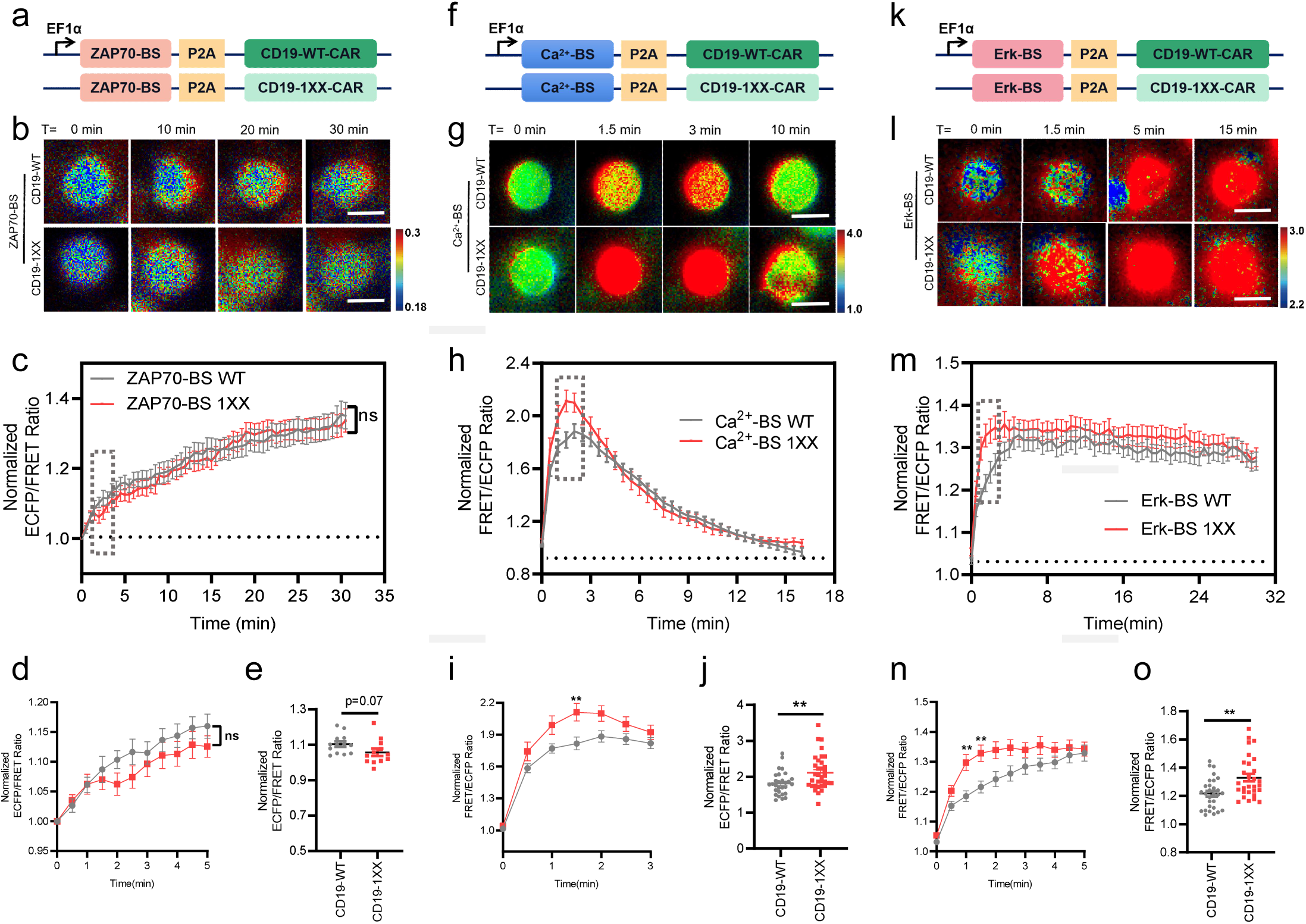
CAR-mediated signaling dynamics in 1XX and WT CAR-T cells revealed by FRET biosensors. (a) Schematic drawings of constructs. The ZAP70-FRET biosensor was co-expressed with WT or 1XX CAR in Jurkat T cells. The CAR-T cells were then dropped onto the 3T3CD19+ cells to monitor the dynamic ZAP70 kinase activations. (b) Representative images of ZAP70 biosensor in T cells after attaching to the 3T3CD19+ cell monolayer. Scale bar,10 μm. (c) Time courses of ECPF/FRET ratio of ZAP70 FRET biosensor in WT or 1XX CAR-T cells (n=13, 13). (d) Time courses of normalized ECFP/FRET ratio of ZAP70 FRET biosensor in WT or 1XX CAR-T cells from 0-5 min (n=13, 13). (e) Normalized ECPF/FRET ratio of ZAP70-FRET biosensor in WT or 1XX CAR-T cells at 2 min (n=13, 13, unpaired two-tailed Student ‘ s t-test, ns, P > 0.05). (f) Schematic drawings of constructs. The Ca2+-FRET biosensor was co-expressed with WT or 1XX CAR in Jurkat T cells. The CAR-T cells were then dropped onto the 3T3CD19+ cells to monitor the dynamic Ca2+ kinase activations. (g) Representative images of Ca2+ biosensor in T cells after attaching to the 3T3CD19+ cell monolayer. Scale bar,10 μ m. (h) Time courses of FRET/ECFP ratio of Ca2+ -FRET biosensor in WT or 1XX CAR-T cells (n=27, 22). (i) Time courses of FRET/ECPF ratio of Ca2+-FRET biosensor in WT or 1XX CAR-T cells from 0-3 min (n=27, 22). (j) FRET/ECFP ratio of Ca2+-FRET biosensor in WT or 1XX CAR-T cells at 1.5 min (n=27, 22, unpaired two-tailed Student ‘ s t-test, **, P < 0.01). (k) Schematic drawings of constructs. The Erk-FRET biosensor was co-expressed with WT or 1XX CAR in Jurkat T cells. The CAR-T cells were then dropped onto the 3T3CD19+ cells to monitor the dynamic Erk kinase activations. (l) Representative images of Erk biosensor in T cells after attaching to the 3T3CD19+ cell monolayer. Scale bar,10 μm. (m) Time courses of normalized FRET/ECPF ratio of Erk-FRET biosensor in WT or 1XX CAR-T cells (n=29, 29). (n) Time courses of normzlied FRET/ECPF ratio of Erk-FRET biosensor in WT or 1XX CAR-T cells from 0-3 min (n=29, 29). (o) Normalized FRET/ECPF ratio of Erk-FRET biosensor in WT or 1XX CAR-T cells at 1.5 min (n=29, 29, **, P < 0.01). All data are mean ± SEM.

## Discussion

1XX design has been shown to be advantageous in the CD19 CAR context against leukemia. The most relevant translational question is whether 1XX modification is also beneficial against solid tumors. Though a prior study reported the 1XX design of a mesothelin-targeting CAR was effective in a pleural mesothelioma mouse model ^[22]^, there was no direct comparison with its WT counterpart and the underlying molecular mechanism were not investigated. Our study demonstrated that 1XX modification can enhance CAR-T cell efficacy against solid tumors in two different CAR settings and tumor models, suggesting this could be a general optimization strategy. The increased efficacy was attributed to the observations that CD19-1XX CAR-T cells extracted from leukemic mice were of higher number and had a higher percentage of CD62^+^CD45RA^-^ central memory T cells than CD19-WT cells ^[15]^. In our study, we tried to isolate CAR-T cells from subcutaneous tumors to analyze their phenotype but we could not detect enough CAR-T cells (data not shown). This was most likely because we injected a low CAR-T cell dose and collected the tumor tissue at the very last time point. We chosed a low CAR-T cell dose as a “stress test” for better comparison of T cells with different CAR constructs ^[20]^. The low number of CAR-T cells in tumors was corroborated by in situ immunostaining of CAR-T cells. Quantitative analysis of immunostaining indicated a trend of more CD19-1XX cells than CD19-WT cells, which was consistent with the results from leukemic mouse model. Nevertheless, we could isolate enough CAR-T cells from mouse spleen instead of tumors for detailed characterization. 1XX mutation in AXL-CAR increased the number of total CAR-T cells by 110.5 fold and these cells displayed increased central memory phenotype and decreased exhaustion trend, which were consistent with the effect of 1XX modification on CD19 CAR.

1XX CAR-T cells have significantly lower cytotoxicity than WT CAR-T cells when killing Panc-1^CD19+^ cells. Previous data showed that CD19-1XX and CD19-WT CAR-T cells display similar cytotoxicity against leukemic NALM6 cells *in vitro* ^[15]^. AXL-specific 1XX CAR-T cells also have decreased killing against A375 tumor cells. Thus it is possible that results of *in vitro* killing assays cannot predict *in vivo* anti-tumor activities of different CAR-T cells. Here 1XX design has already been combined with the TRAC-targeting of CAR to take advantage of two optimization methods ^[19]^. Whether 1XX can be combined with other known therapeutic enhancement strategies, such as IL-12, PD-1 blocking scFv or IL-7/CCL19, to further enhance CAR-T performance against solid tumors, warrants further investigation.

RNAseq analysis of CD19-1XX and CD19-WT cells revealed their global transcriptional difference 24h after target recognition ^[15]^, but how 1XX mutation leads to differential gene expression was still elusive. Here we employed different FRET biosensors to probe the signaling dynamics of three key molecules downstream CAR activation (Fig. 6). Each ITAM from CD3ζ has two tyrosines, both of which need to be phosphorylated to recruit ZAP70 so that ZAP70 can be phosphorylated by Lck ^[9]^. But singly-phosphorylated ITAMs was shown to instead recruit Shp1 phosphatase, which in turn inhibits Lck activity through negative feedback ^[23]^ (Fig. 6). Thus 1XX with reduced ITAMs could enhance Lck activity by decreasing the amount of singly-phosphorylated ITAMs and associated Shp1. The total ZAP70 activity of 1XX would be the net result of these two opposing effects, as reduced ITAM numbers decreased the amout of bound ZAP70 but increased the activation of bound ZAP70 by enhanced Lck activity. Our FRET data showed that even though ZAP70 activity in 1XX was not statistically different from that in WT, the trend was lower (Fig. 5e).

**Fig.6:**
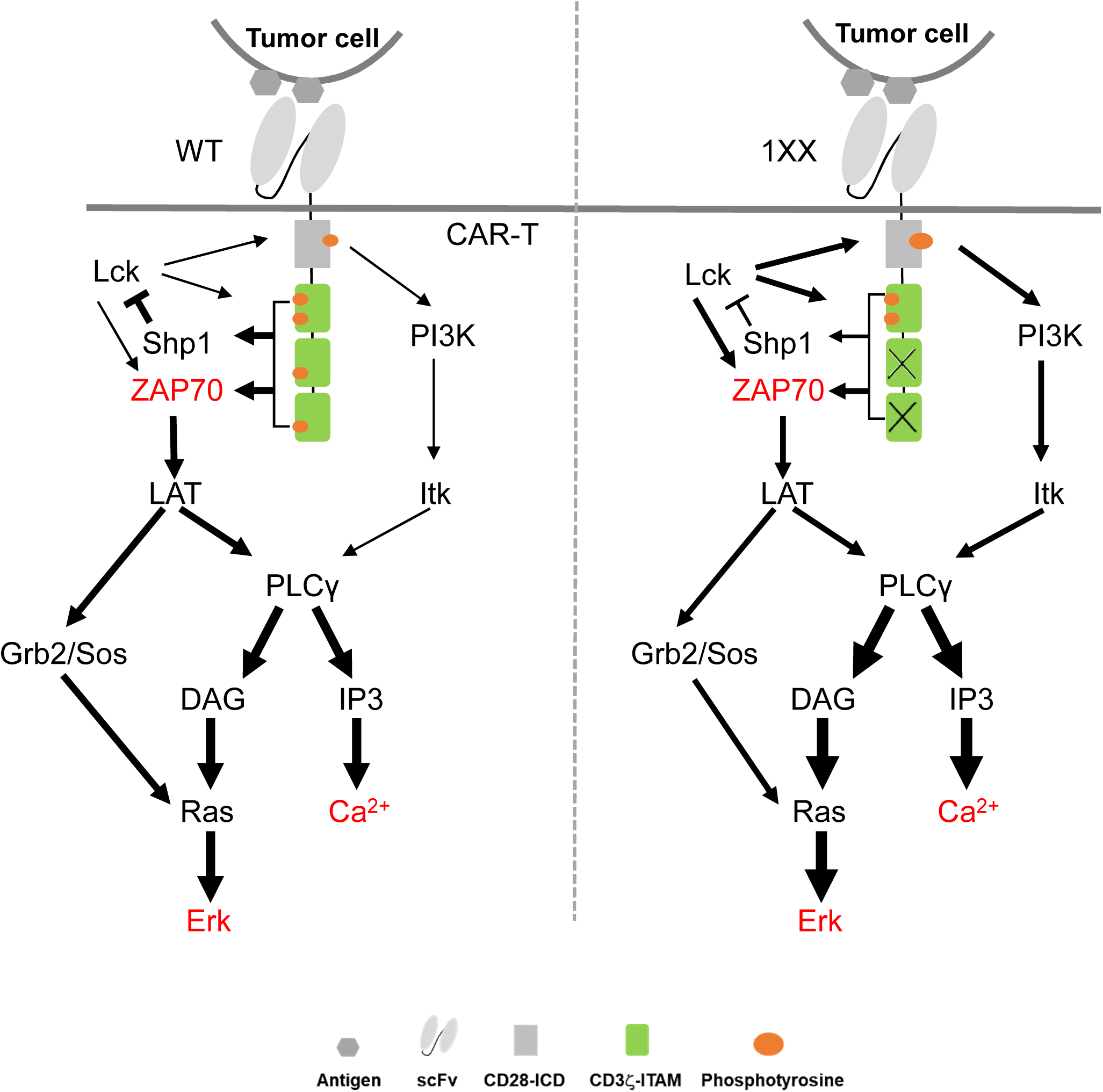
Signaling dynamics of three key molecules downstream WT (left) and 1XX (right) CAR activation. Upon CAR binding to antigen, Lck phosphorylates CD28 and CD3 ζ signaling domains. Doubly-phosphorylated ITAMs recruit ZAP70 and ZAP70 gets activated by Lck. Singly-phorphorylated ITAMs recruit Shp1 to inhibit Lck by negative feedback. Activated ZAP70 phosphorylates LAT and subsequently activates both Ca2+ and Erk. Phosphoryalted CD28 also activates Ca2+ and Erk through PI3K pathway. In 1XX, reduced ITAMs lead to decreased Shp1 inhibition and enhances Lck activity. Thus PI3K-dependent Ca2+ and Erk pathway gets strenghtend. Net ZAP70 activity remains unaltered from two opposing effects: decreased ZAP70 binding from reduced ITAMs and increased ZAP70 phosphorylation by Lck.

If only CD3ζ-ZAP70 pathway was considered, 1XX mutation should not aleter the Ca^2+^ and Erk activation through LAT. But in the context of CD28-based second generation CAR, CD28 cytosolic domain after being phosphorylated by Lck can also augument Ca^2+^ and Erk signaling through PI3K activation ^[11,12]^ (Fig. 6). As 1XX can enhance Lck activity through reduced singly-phosphorylated ITAMs and associated Shp1 ^[24,25]^, CD28 phosphorylation by Lck in 1XX CAR may be strengthened, which promotes higher or faster PI3K activation ^[11,12]^ (Fig. 6). Our FRET-based imaging revealed that 1XX resulted in significantly higher Ca^2+^ evelation and more rapid Erk activation than WT upon antigen binding (Fig. 5). Increased Ca^2+^ would bind to calcinurin and activate transcription factors of the NFAT family to regulate cytokine gene expression and other genes critical for the T cell functions ^[26]^. Erk could further activate transcription factors, such as Elk-1 and AP-1 ^[27]^. Particularly, Erk pathway was shown to be pivotal for both human memory Th17 cells and CD8 T cell differentiation ^[28-30]^, suggesting that faster Erk activation kinetics of 1XX may be linked to CAR-T cell differentiation.

## Conclusions

Our study established the superior anti-tumor effect of 1XX CAR in two preclinical solid tumor models and revealed the delicate changes of signaling dynamics caused 1XX CAR, paving the way for the clinical translation of 1XX CAR in treating solid tumors.

## Methods

### Cells lines and culture conditions

Panc-1 cells were transduced to express human CD19 as described previously ^[20,31]^, and Panc-1^CD19+^ cells were sorted by Flow Cytometry. Panc-1^CD19+^ and A375 cells were transduced to express firefly luciferase (FFLuc)-2A-green fluorescent protein (GFP), and GFP^+^ cells were sorted by Flow Cytometry. Panc-1, A375, 293T and NIH/3T3 were cultured in Dulbecco’s modified Eagle medium (DMEM, Gibco) medium with 10% FBS (Vistech) and 1% penicillin/streptomycin(Gbico). Jurkat cell was cultured in RPMI-1640 medium (Gibco) with 10% FBS and 1% penicillin/streptomycin.

### RNP production

RNPs were produced by complexing gRNA and Cas9 protein. Modified guide RNAs (gRNAs) was synthesized by GeneScript. Guide RNAs were reconstituted at 1μg/μL in RNase free water, Cas9 proteins were produced by our lab as described in our previous work ^[32]^, In brief, they were complexed in 2:1 gRNA to Cas9 molar ratio at room temperature for 20 min then electroporated into T cells immediately after complexing ^[32]^.

### Plasmid construction

Based on pAAV-TRAC-1928z plasmid (as described in our previous work ^[32]^, named it as CD19-WT) we designed and cloned the pAAV-TRAC-1928z-1XX (named as CD19-1XX). Briefly, the CD19 CAR comprises a single chain variable fragment ^[19]^ scFv specific for the human CD19 (AXL CAR comprises a single chain variable fragment of anti-AXL ^[33]^ specific binding to human AXL), preceded by a CD8a leader peptide and followed by CD28 hinge, transmembrane and intracellular regions and CD3ζ intracellular domain. The CAR cDNA is followed by the bovine growth hormone polyA signal (bGHpA). In briefly, pAAV-TRAC-AXL28z/1XX plasmids were constructed with following primers. AAV-AXL-forward primer: GTGGAGGAGAATCCCGGCCCCatggctctcccagtgactgccctactg; AAV-AXL-reverse primer: GCAACTAGAAGGCACAGTCGcctagggatttagcgagggggcagggcctg, the anti-AXL scFv fragment was obtained by PCR and insert into pAAV-TRAC-1928z-WT or pAAV-TRAC-1928z-1XX plasmid which was digested out 1928z-WT or 1928z-1XX fragment by NcoI/AvrII, respectively. The pAAV-TRAC-AXL28z/1XX plasmids were prepared using standard molecular biology techniques as described previously ^[15]^. The Lenti-ZAP70-BS-CD19 CAR, Lenti-Ca^2+^-BS-CD19 CAR and Lenti-Erk-BS-CD19 CAR plasmids were constructed as in our previous work ^[16]^ for FRET analysis.

### Isolation and expansion of human T cells

Human PBMCs were isolated from peripheral blood of healthy volunteers. Ethical permission was granted by the School of Medicine, Zhejiang University. All blood samples were handled following the required ethical and safety procedures. Peripheral blood mononuclear cells were isolated by ficoll density gradient centrifugation (Dakewe) and T cells were purified using the Pan T Cell Isolation Kit (Miltenyi Biotec), and stimulated with CD3/CD28 T cell Activator Dynabeads (Thermofisher) as described previously ^[19]^ then cultured in X-VIVO 15 Serum-free Hematopoietic Cell Medium (Lonza), which supplemented with 10% fetal bovine serum (Vistech), 5ng/mL IL7 (interleukin-7) and 5ng/ml IL15 (interleukin-15) (Novoprotein) for experiments. The medium was changed every 2 day, and cells were plated at 10^6^ cells/mL. After stimulated for 48 h, human T cells were debeaded for gene targeting experiments.

### CAR-T cell production

RNPs was electroporated into T cells 2 days after CD3/CD28 bead stimulation. Before electroporation, debeaded T cells were resuspended in X-VIVO 15 medium. For each reaction, 3×10^6^ cells were mixed with 3.6 μL (80 pmol) RNPs in a total volume of 120 μL and transferred to a 120ul cuvette and electroporated with an Cell Electroporator (CELETRIX). 30 min later, AAV virus (MOI=1×10^5^) (Vigene Biosciences, China) were added. Following electroporation cells were immediately transferred into culture medium (X-VIVO 15 with 10% FBS and 1% penicillin/streptomycin). 2 h later, IL-7(5 ng/mL) and IL-15(5 ng/mL) were added. Then medium was changed every 48 h. 7 days after electroporation, cells were harvested for fluorescence-activated cell sorting (FACS) analysis to determine the knock-in efficiency in each condition.

### ELISA assay

Briefly, ELISA plates were coated with 100 μL/well of 2 μg/mL AXL-ECD protein. DAXL-88 was diluted, added to the wells at 15, 5, 1.67, 0.56, 0.187, 0.062, 0.02, 0.007, 0.002 and 0.0008 μg/mL, the data was measured with an ELISA reader as described in our previous work ^[33]^.

### Flow cytometry

The following fluorophore-conjugated antibodies were used. For CD19 CAR staining, an Alexa Fluor 647 AffiniPure F(ab’)2 Fragment Goat Anti-Mouse IgG was used (Jackson ImmunoResearch); for AXLCAR staining, an Alexa Fluor 647 AffiniPure anti-HA tag Goat Anti-Mouse IgG was used (BD Bioscience). A375 stained with anti-AXL PE-conjugated antibodies (FAB154P, R&D); and the DAXL-88 (anti-AXL) binding assay used Alexa Fluor 647 AffiniPure anti-Fc tag Goat Anti-Human IgG (410711, BioLegend). Anti-Human CD4 antibody (BD, Horizon, BUV395), anti-Human CD8 antibody (BD, Pharmin, APC-CY™7), anti-Human CD279(PD-1) antibody (Invitrogen, eBioscience, PE-Cyanine7), anti-Human CD223(LAG-3) antibody (Invitrogen, eBioscience,, Percp-efluor™710), anti-Human CD45RA antibody (Invitrogen, eBioscience, FITC), anti-Human CD62L antibody (Invitrogen, eBioscience, efluor450). For cell counting, CountBright Absolute Counting Beads were added (Invitrogen) according to the manufacturer’s instructions. For in vivo experiments, Fc receptors were blocked using FcR Blocking Reagent, mouse (Miltenyi Biotec). CytoFLEX (Beckman) was used with FlowJo 7.6 software for analysis. All FACS plots presenting CAR-T cell phenotype data were conducted on gated APC positive CAR-T cells.

### Antigen stimulation and cytokines analysis assay

Nine days after gene targeting, the CAR-T cells and irradiated target cells NIH/3T3-CD19 or A375 were cocultured at an E:T ratio of 4:1 without the addition of exogenous cytokines for 24 h, then cell culture supernatant were harvested and analysed using a BD Cytometric Bead Array (CBA) Human Soluble Protein Master Buffer Kit (BD Biosciences) according to the manufacturer’s instructions. The detection reagent is a mixture of phycoerythrin (PE)-conjugated antibodies, which provides a fluorescent signal in proportion to the amount of bound analyte. Soluble cytokine can be measured using flow cytometry to identify particles with fluorescence characteristics of both the bead and the detector.

### Proliferation assay

For the proliferation assay, CAR-T cells were labeled with Carboxyfluorescein diacetate succinimidyl ester (CFSE) using a kit from Beyotime. Briefly, cells were resuspended in RPMI 1640 with 10% FBS at a final concentration of 5×10^6^ cells/mL, and CFSE solution was added at the suggested working concentration. The CAR-T cells were incubated at 37 °C for 10 min, and then washed three times with RPMI 1640 with 10% FBS. CFSE-labeled cells were futher plated in 24-well plates, cocultured with irradiated 3T3^CD19+^ or A375 cells (E:T=4:1) for 72 h. Unstimulated but CFSE-labeled CAR-T cells served as control. 72h later CAR-T cells stained with CFSE were analyzed by CytoFLEX (Beckman) with FlowJo 7.6 software.

### Cytotoxicity assay

Nine days after gene targeting, CAR-T cells were collected for luciferase-based cytotoxicity assay using FFluc-GFP Panc-1^CD19+^ or A375 as target cells. The target (T) and effector (E) cells were co-cultured in triplicates at indicated E/T ratios using black 96-well flat plates with 2.5×10^4^ target cells in a total volume of 100 μL per well in X-vivo 15 medium. Target cells alone were plated at the same cell density to determine the maximal luciferase expression (relative light units (RLU)); 18 h later, 100 μL luciferase substrate (Goldbio) was directly added to each well. Emitted light was detected in a luminescence plate reader. Lysis percentage was determined as (1−(RLUsample)/(RLUmax)) × 100.

### Immunofluorescence of tumor tissues

The mice were euthanized by carbon dioxide (CO_2_) inhalation, and the tumors were harvested and fixed in 4% paraformaldehyde. Tumor tissue were sectioned for anti-CD4 (GB13064-1, Servicebio) and anti-CD8 (GB13068, Servicebio) staining. All procedures followed the manufacturer’s protocol. In brief, tumor tissue slides were incubated at 65°C for 1h, blocked with PBS containing 3% BSA for 30 min at room temperature, and then incubated with primary antibody at 4°C overnight. Secondary antibody was incubated at room temperature for 50 min in dark condition. Then slides were incubate with DAPI solution at room temperature for 10 min in dark condition. Images were acquired by fluorescent microscopy. Nucleus is blue by labeling with DAPI. Positive cells are green or red according to the fluorescent labels used.

### Live cell imaging with FRET-based biosensors

The CD19 CAR-T cells expressing biosensors were generated by lentivirus transduction of Jurkat T cells ^[16]^. For imaging of CAR signaling upon antigen stimulation, CAR-T cells were dropped on the glass-bottom dishes that have been coated with the NIH-3T3 cells expressing CD19 antigens. The time-lapse fluorescence images were taken with a Nikon Eclipse Ti inverted microscope at an interval of 30 s. The W-VIEW GEMINI imaging splitting optics (Hamamatsu, Japan) with an iXon Ultra 897 camera was used to capture the ECFP (a 474/40 nm emission filter) and FRET (a 535/25 nm emission filter) fluorescent signals simultaneously. During imaging, the cells were maintained with 5% CO_2_ at 37 °C using the Tokai Hit ST Series Stage Top Incubator (Tokai Hit, Japan).

### Mouse systemic tumor model

6 to 8 weeks old NOD/SCID/IL-2Rγ null (NSG) mice (Shanghai Jihui, China) were used. All mice were housed at the Westlake University under pathogen-free conditions, and all procedures were approved by the ethical committee of Westlake University. Mice were subcutaneously (s.c.) injected a total of 1×10^6^ target tumor cells (Panc-1^CD19+^ or A375) into the right flanks. 7 days later 1×10^6^ CD19-specific or 2.5×10^6^ AXL-specific CAR-T cells were injected. Tumor size was recorded every 3-4 days using a vernier caliper. Tumor volume was calculated as follows: tumor volume = long diameter × (short diameter^2^)/2. NSG mice were euthanized when the tumor size exceeded 14mm (Panc-1^CD19+^ model) or 19mm (A375 model), so Panc-1^CD19+^ model mice were euthanized and tissue was harvested 53 days after CAR-T cell injection, A375 model mice were euthanized and tissue was harvested 26 days after CAR-T cell injection. For each group, 3-5 individual mice were included. Stratified randomization based on initial tumor size was used. No animal data were excluded in this study. No strategy was used to minimize potential confounders. D.Y and J.C. were aware of the group allocation at the different stages of the experiment.

### Statistical analysis

Student *t* test or multiple *t* test was carried out using GraphPad Prism version 7.0 (GraphPad Software Inc). ns, P > 0.05; *, P < 0.05; **, P < 0.01; ***, P < 0.001 and ****, P < 0.0001.

## Abbreviations

ITAMs: Immunorecpetor tyrosine-based activation motifs
CAR-T: Chimeric antigen receptor engineered T cells
WT: Wild type
CFSE: Carboxyfluorescein diacetate succinimidyl ester
IFN: Interferon
IL: Interleukin
TNF: Tumor necrosis factor
FRET: Fluorescence resonance energy transfer
ZAP70: 70-kDa zeta-associated protein
Ca^2+^: Calcium
Erk: Extracellular regulated protein kinases
FBS: Fetal bovine serum
DMEM: Dulbecco’s Modified Eagle Medium
RPMI: Roswell Park Memorial Institute

## Acknowledement

The authors thank the support of Zhejiang Provincial Key Laboratory of Immunity and Inflammatory diseases. The authors thank Zhejiang University School of Medicine core facilities for their support.

## Authors’ contributions

Y.D. and J.S. designed the studies and conceived the experiments. Y.D., J.C. and XH.M. performed most of the experiments; L.L., K.S., X.W., Y.W., Z.H. and H.L. conducted data analysis; Y.D. and J.S. wrote the manuscript. Y.H., C.Z., X.G., Y.W. and M.S. discussed the manuscript, X.G., J.S. were responsible for funding acquisition. All authors read and approved the final manuscript.

## Availability of data and materials

All data obtained and/or analyzed during the current study are available from the corresponding authors on reasonable request.

## Ethics approval and consent to participate

All animal studies were reviewed and approved by the Ethical Committee of Westlake University.

## Consent for publication

All authors have given consent for the publication of this manuscript.

